# A brief exposure to rotenone alters microtubule dynamics resulting in aberrant elongation of primary cilia with impaired function

**DOI:** 10.64898/2025.12.29.696880

**Authors:** Priyadarshini Halder, Rupsa Mondal, Eshona Chakraborty, Shubhra Majumder

## Abstract

Rotenone, an environmental toxin, is widely used to model Parkinson’s disease owing to its well-studied effect to inhibit mitochondrial complex I function and to increase Reactive Oxygen Species (ROS), thereby leading to dopaminergic neuron degeneration. Less appreciated is its impact on microtubules (MT), except for suggesting that it promotes microtubule depolymerization, raising the question of whether rotenone also impacts primary cilia (PC) that are microtubule-based signalling hubs, which are essential for neuronal function.

Using hTERT-RPE1 (or RPE1) cells, which assemble PC upon serum starvation, we discovered that brief exposure to rotenone (2-6 h) at low concentration (100 nM) causes a striking elongation of PC and hyperacetylation of cytoskeletal microtubules. At this concentration, rotenone does not significantly alter mitochondrial dynamics or bioenergetics, and these phenotypes could not be reversed by pre-incubating cells with ROS scavenger NAC, or supplementing cells with NR. Thus, these observed effects of rotenone treatment in quiescent cells are predominantly independent of mitochondrial toxicity. Depletion of α-TAT1, the α-tubulin acetyltransferase, eliminated microtubule acetylation without preventing ciliary elongation, suggesting that rotenone drives PC extension through additional mechanisms. Importantly, rotenone increases soluble tubulin pools, owing to its microtubule destabilizing effect, which are likely the mechanism contributing to ciliary axoneme elongation as well as hyperacetylation of cytoskeletal microtubules. Critically, such aberrant increase in PC length impaired Sonic Hedgehog (SHH) signalling, which is exclusively transduced by PC during embryonic development and is critical for maintaining adult tissue homeostasis. Thus, our findings reveal that even brief, low-dose rotenone exposure induces aberrant elongation of PC in quiescent cells that assemble PC, while simultaneously disrupting the function of PC.

Based on our observation and previous studies, we propose that such ciliary dysfunction represents an underappreciated mechanism by which rotenone contributes to the pathogenesis of neurodegenerative disease via affecting neuronal primary cilia.

## Introduction

Environmental toxins have long been implicated as significant contributors to the development of neurodegenerative diseases, including Parkinson’s disease (PD)[1]. Among these, rotenone, a naturally derived isoflavonoid compound commonly used as a pesticide and insecticide, has been extensively studied as a reliable experimental tool for modelling PD both in cell culture and *in vivo* mouse or rate model [1, 2]. Rotenone is a well-known inhibitor of mitochondrial complex I (NADH: ubiquinone oxidoreductase) of the electron transport chain, thereby may impair oxidative phosphorylation and ATP production. Also, rotenone treatment leads to increased production of reactive oxygen species (ROS), DNA damage and mitochondrial dysfunction, all of which may ultimately contribute to cell death or degeneration of dopaminergic (DA) neurons, a hallmark of PD pathogenesis [3]. Separately, exposure to rotenone may also enhance age-dependent expression of cell death inducing protein Sarm1 and associated inflammatory responses, resulting in the dopaminergic neurotoxicity[4]. However, few studies also hinted that rotenone-induced neurotoxicity arises due to its effect on microtubular dynamics and may not be due to inhibition of mitochondrial complex I or altered mitochondrial morphology[2, 5]. In fact, rotenone binds to microtubules, and promotes microtubule depolymerization[6]. This is particularly important as exposure to low dose of rotenone may not be sufficient to cause cell death but may affect cellular physiology and related phenotypes, often seen in DA neurons during PD pathology[7]. Based on this information we wondered if brief exposure to rotenone affects structure and function of primary cilia that are solitary, non-motile organelle extending from the surface of most post-mitotic mammalian cells.

The primary cilium (PC) contains microtubule-based axoneme that is assembled on basal body and ensheathed under ciliary membrane, which is continuous with the cell membrane[8]. PCs perform sensory functions and serve as critical signalling hub that coordinates multiple developmental and homeostatic pathways, including the Sonic Hedgehog (SHH), Wnt, PDGF, Notch, and TGF-β signalling pathways[9–11]. Basal bodies are transformed mother centriole, the oldest of the centriole pair in a post-mitotic centrosome. In cycling cells, the assembly and disassembly of the PC are temporally linked to the cell cycle and centriole duplication cycle[12], cytoskeletal dynamics, and post-translational modifications on tubulin, such as acetylation, glutamylation [13]. Dysfunction of PC or motile types of cilia, conditions broadly referred to as ciliopathy, is associated with a wide range of human diseases, including polycystic kidney disease, Bardet-Biedl syndrome, Joubert syndrome, retinal degeneration, obesity, and neurodevelopmental disorders[14]. PCs in neurons play crucial roles in neural development, neuronal differentiation, synaptic integration, and maintaining tissue homeostasis. Importantly, defects in ciliary structures or signalling have been increasingly linked to neurodegeneration, with growing evidence supporting the role of PC dysfunction in disorders such as Parkinson’s and Alzheimer’s diseases[15–17]. Neurons rely on ciliary signalling to accurately interpret extracellular cues to maintain cellular health. Thus, even subtle alterations in PC length and structure may affect the signalling competency, with profound consequences[18].

Thus, it is important to mechanistically investigate the effects of environmental neurotoxins, such as rotenone, on PC and downstream signaling to understand the associated pathology. However, there are only limited studies in this aspect, which either used a longer duration of significantly higher concentration of rotenone than that is required to inhibit mitochondrial complex I, and did not consider the possible effect of rotenone exposure on microtubule dynamics[19–21]. Also, in some cases, the condition and the cell types were not ideal for robust PC assembly. In cell culture model, human hTERT-RPE1 (or RPE1) cells or mouse NIH3T3 fibroblasts are used to study mechanistic details of PC assembly-disassembly, since these cells, upon serum starvation (24-48 h), are arrested in G0-phase or cellular quiescence, and therefore 80-90% of the cells assemble PC, while only 15-18% of asynchronously growing RPE1/3T3 cells assemble PC in presence of serum. Also, upon serum addition, these cells synchronously disassemble PC, thereby reducing ciliation to 40-45% within 20 h.

Here, we discovered that microtubule destabilization due to brief exposure to rotenone induced hyperacetylation of cytoplasmic and ciliary microtubules and striking elongation of PC in quiescent RPE1 cells. Our study clearly indicated that such aberration in ciliary length is not due to increase in cellular ROS or loss of mitochondrial bioenergetics. Importantly, we further demonstrated that rotenone-induced elongated cilia are functionally impaired, as evidenced by disrupted SHH signaling. These findings reveal an unexpected and underappreciated mechanism of rotenone toxicity, which may likely be seen in quiescent, post-mitotic dopaminergic neurons, with pathological implications.

## Results

### Brief exposure to rotenone induces hyperacetylation of cytoskeletal microtubules and PC elongation

Recently in a collaborative study, we demonstrated that rotenone treatment, although at a higher concentration (5 μM) and longer duration (24 h), led to hyperacetylation of cytoplasmic microtubules in asynchronous population of neuroblastoma cells resulting in G2-M arrest[22]. In the present study, we aimed to examine how brief, low-dose rotenone exposure alters the microtubule dynamics and primary cilia structure and function in quiescent cells that already assembled PC. Therefore, we sought to conduct our study in 24 h serum-starved RPE1 cells that are non-transformed diploid cells. Accordingly, these cells were serum-starved for 20 h when more than 85% of these cells contain PC, followed by treating with rotenone or DMSO, the solvent control (Fig.1A). To determine a suitable concentration and treatment duration of rotenone for studying the early cellular effects of rotenone on microtubule dynamics and PC, we first treated quiescent RPE1 cells with increasing durations of rotenone at a final concertation of 100 nM. We stained these cells using antibodies against acetylated α-tubulin (Ac-tub) that marks both cytoplasmic and axonemal acetylated microtubules and Arl13B, a small GTPase that is exclusively localize to ciliary membrane. We observed drastic increase in acetylation of cytoplasmic microtubules within 1 h of rotenone treatment, in comparison to control. DMSO treatment that showed basal acetylation of cytoplasmic microtubules and hyperacetylation of ciliary axoneme (Fig.1B). Such increase in microtubule acetylation could also be determined by immunoblotting the whole cell extract, where the level of α-tubulin remained unaltered after rotenone treatment (Fig.1C). Importantly, rotenone treatment led to a significant increase in PC length as judged by Arl13B staining even at 1 h, which became more pronounced at 2 h and 6 h (Fig. 1B). We did not observe any noticeable change in cell morphology or cell death upon rotenone treatment for this tenure. In fact, even after 24 h of treatment, cell viability remains almost 95% when these quiescent RPE1 cells were treated at a concentration of 100 nM (Fig.1D). Although we were looking for the shortest duration of rotenone exposure, for the sake of all the mechanistic assays of this study associated with aberrant elongation of PC, we chose 4 h exposure to rotenone at 100 nM as the optimal treatment condition balancing robust cellular and ciliary phenotypes with minimal loss of cell viability. Under this selected condition, rotenone treatment produced striking cytoskeletal microtubular rearrangements and markedly elongated PC relative to DMSO-treated cells as revealed by confocal imaging (Fig. 1E).

**Figure 1.**
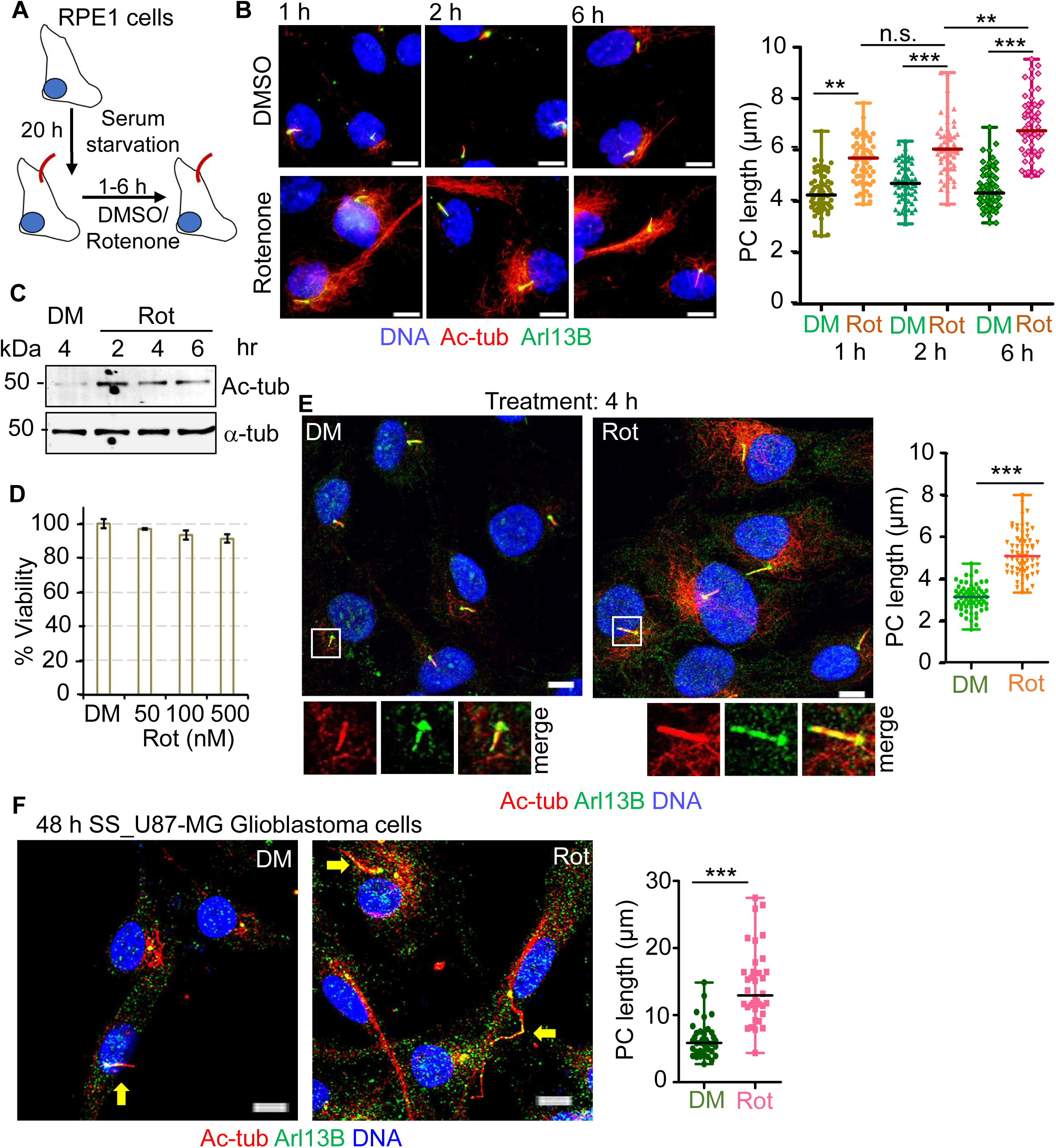
Short-term, low-dose rotenone induces rapid primary cilia elongation and microtubule hyperacetylation without compromising cell viability. (A) Schematic of the experimental workflow using hTERT-RPE1 (or RPE1) cells. (B) Representative immunofluorescence images of serum-starved RPE1 cells treated with DMSO or 100 nM rotenone for 1, 2, or 6 h and stained for acetylated α-tubulin (Ac-tub) and Arl13B. Here and in other figures, Hoeschst33358 is used to stain DNA (blue) and scale bar is 5 µm. Here and for all other experiments, the length of PC (in µm) is determined from at least 100 randomly picked cells per experimental replicate and are presented in a dot-plot, where the marker indicates the median, and the whiskers represent minimum and maximum values for each series. Measurement of ciliary length shows a significant increase in length over time following rotenone treatment compared with DMSO controls. Here and in all other cases, p value was determined by unpaired t-test, where *** indicates p<0.001, ** indicates p < 0.01, * indicates p < 0.05 and n.s. indicates non-significant. (C) Immunoblot analysis of acetylated α-tubulin in RPE1 cells treated with 100 nM rotenone for 2, 4, 6 h and compared with DMSO control. α-tubulin was used as a loading control. Rotenone causes a clear increase in Ac-tub levels in treated cells, while total α-tubulin remains unchanged. (D) Cell viability assessed by MTT assay of cells after 4 h treatment with increasing concentrations of rotenone. Bars represent % viability (compared to solvent control) of indicated samples where values represent mean ± S.D., n=3. (E) Representative confocal images showing cytoplasmic microtubule acetylation in serum-starved RPE1 cells treated with DMSO or 100 nM rotenone for 4 h, while the dot plots show significant elongation of PC length in rotenone treated cells compared to control. (F) Representative images of serum-starved U-87 MG cells treated with DMSO or 100 nM rotenone for 4 h and stained for indicated ciliary markers, with the dot plots of PC length quantification of cilia length demonstrating significant elongation upon rotenone treatment.

Next, to determine whether this effect is conserved in cells originated from nervous system we examined human U87-MG glioblastoma cells. Only 5-8% of these cells are known to assemble PC under 24-48 h serum starvation [23]. Like RPE1 cells, 4 h treatment of rotenone induced pronounced PC elongation in U87-MG cells (Fig. 1F), which mostly were more than twice the length of the PC observed in control cells. Expectedly, cytoplasmic microtubules were hyperacetylated upon rotenone treatment compared to DMSO treatment (Fig. 1F).

Collectively, these results demonstrate that brief, low-dose rotenone exposure is sufficient to induce hyperacetylation of cytoskeletal microtubules likely via altering cytoskeletal microtubular dynamics. Also, such treatment induces drastic PC elongation in quiescent cells that supports PC assembly and also in cancer-derived cells that rarely have PC. Since cellular health and viability remained unaltered under this treatment condition, rotenone-mediated mitochondrial toxicity might not operate during this treatment. Therefore, it is possible that altered microtubule dynamics and ciliary remodelling represent early events in rotenone-mediated loss of cellular physiology, often associated with rotenone-induced neuronal degeneration.

### Rotenone-induced elongation of PC is accompanied by impaired SHH signalling

Because PCs function as essential signalling organelles, we next asked whether the rotenone-induced elongated PCs remain similarly competent to transduce signals. In particular, we checked SHH signalling pathway that is based on differential processing of Gli transcription factors within PC, which ultimately results in upregulation of the SHH-target genes. We first utilized the recently developed qRT-PCR based assay to assess the expression of canonical SHH target genes GLI1, PTCH1, and HHIP when SHH signaling was initiated by incubating serum starved cells with Smoothened Agonist for 20 h (SAG, Fig.2A). When SAG-treated cells were treated with DMSO for the last 4 h, an expected increase in the expression of the three SHH target genes compared to the SAG-untreated cells were observed. When the SAG-treated cells were exposed to rotenone for the last 4 h, PC length increased significantly compared to DMSO treatment (Fig.2B). However, no further increase in the expression of those genes were observed, albeit such increase in average PC length in rotenone-treated cells (Fig.2C). This result indicates that increase in PC length does not enhance its ability to traduce SHH signaling likely due to some aberration in these longer PCs. Because Smoothened (Smo) trafficking into and along the ciliary axoneme is a hallmark of SHH pathway activation, we next examined Smo localisation along PC in SAG-treated cells in presence of rotenone or DMSO. For this purpose, we incubated the serum starved RPE1 cells with SAG for 6 h, along with rotenone or DMSO for the last 4 h, to observe ciliary localization of Smo during the early phase of mimicking SHH signaling. In DMSO-treated cells, Smo was readily detectable along the ciliary axoneme in most PCs, consistent with robust pathway activation (Fig. 2D). Strikingly, in rotenone-treated cells, Smo failed to localise along the length of PCs in most cells, suggesting defective trafficking of Smo that may result aberrant SHH signaling (Fig.2D). Indeed, quantitative analyses show a significant decrease in total Smo fluorescence intensity on the PCs marked by Arl13B, upon rotenone treatment compared to DMSO treatment of SAG-activated cells (Fig.2D). Together, these findings demonstrate that elongated PCs generated under rotenone exposure present are aberrantly functional in response to SHH stimulation, likely due to core structural aberration of PCs. This also raises the possibility that such structural defects in PC perhaps not arise from rotenone-induced impairment of mitochondrial function, but from a distinct mechanism.

**Figure 2.**
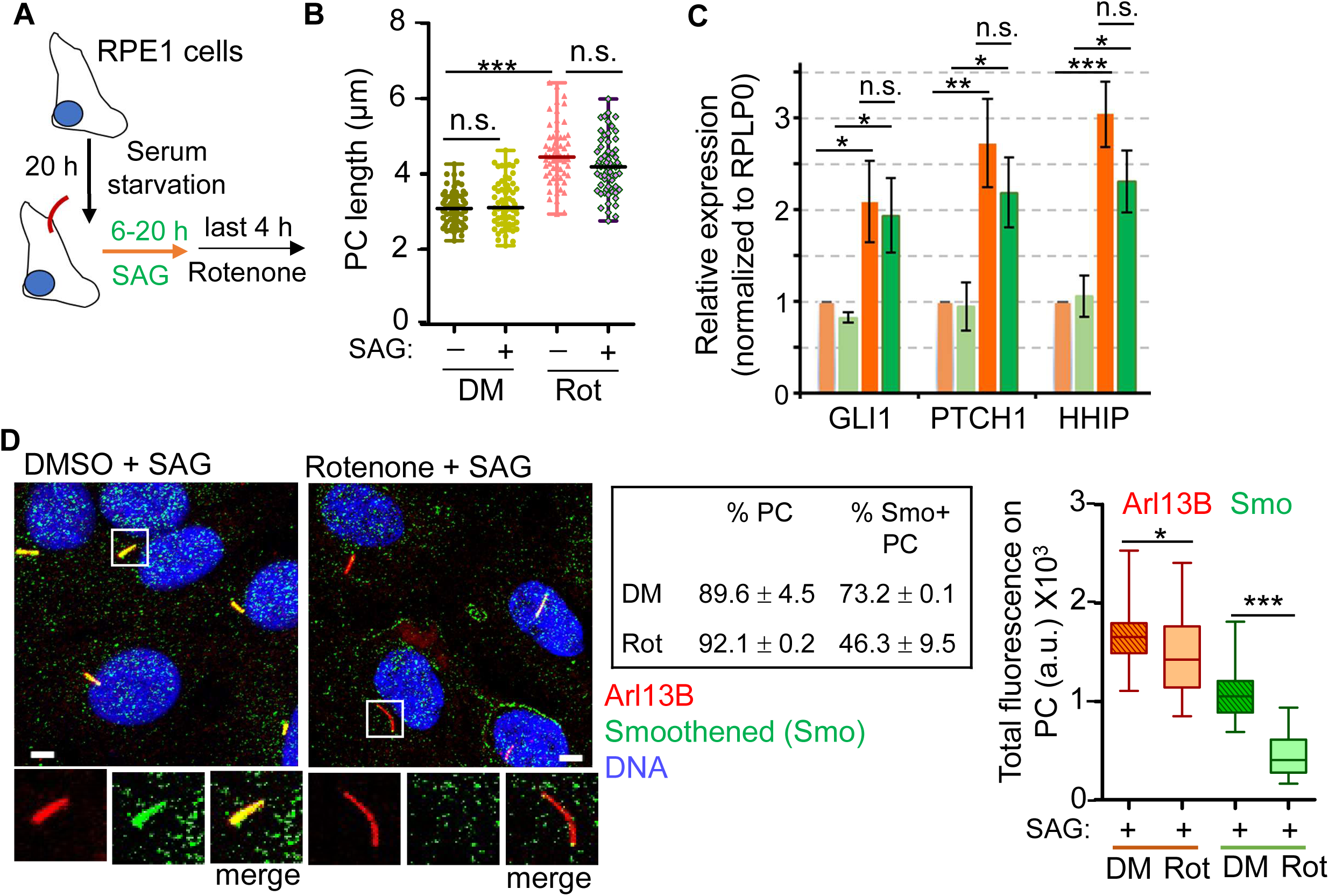
Elongated cilia induced by rotenone are functionally impaired for Sonic Hedgehog signalling. (A) Schematic to examine the canonical Sonic Hedgehog (SHH) signalling pathway. (B) Cells were treated with DMSO or rotenone in presence or absence of SAG, and stained for ciliary markers. The PC length of these cells is shown as dot plot. (C) Quantitative real-time PCR analysis of canonical SHH target genes (GLI1, PTCH1, HHIP) in serum-starved RPE1 cells treated with DMSO or 100 nM rotenone, with or without SAG stimulation. The relative expression (2^-ΔΔCT^) of β-ACTIN normalized transcript levels of the indicated genes in serum-starved RPE1 cells compared to the control treatment (DMSO, no SAG) are plotted as bars, where values represent mean ± S.D., n=3. (D) Rotenone disrupts Smo trafficking into the cilium. Representative confocal images show Smo localisation in DMSO- and rotenone-treated cells following SAG stimulation, with cilia marked by Arl13B. % of total cells containing Arl13B-positive PC are shown in the table. Background-corrected total intensity of Smo signal along PC in different samples are presented in a box and whisker diagram, where boxes indicate lower and upper quartiles, the marker in the box (–) indicates the median, and the whiskers represent minimum and maximum values for each series. 80-100 cells with PC were analyzed for each sample.

### Rotenone-induced PC elongation and altered microtubule hyperacetylation in quiescent cells occur independently of its mitochondrial toxicity

Rotenone inhibits mitochondrial respiratory chain complex I at 10-100 nM concentration in a variety of cells, and also generates Reactive Oxygen Species (ROS), which are considered the major causes of rotenone-induced cell death. Usually, the later requires higher concentration of rotenone (1-5 μM) and 12-24 h of treatment[3, 24]. Now, we asked whether the effect of rotenone on microtubule modification and axoneme elongation arise from impaired mitochondrial function. We first observed that the mitochondrial distribution and size as judged by MitoTracker Red incorporation and staining did not change significantly in rotenone treated serum starved RPE1 cells, compared to DMSO treatment (Fig.3A). Next, we examined mitochondrial dynamics and membrane potential using MitoTracker Green in live cells during the last 30 min of acute treatment with 100 nM rotenone or DMSO. Since there was no appreciable difference in MitoTracker Green intensity or mitochondrial morphology or abundance between the two samples, we conclude that mitochondrial membrane potential or mitochondrial dynamics remain largely nonperturbed upon the brief treatment of rotenone, which causes microtubule hyperacetylation and aberrant PC elongation (Fig. 3B).

**Figure 3.**
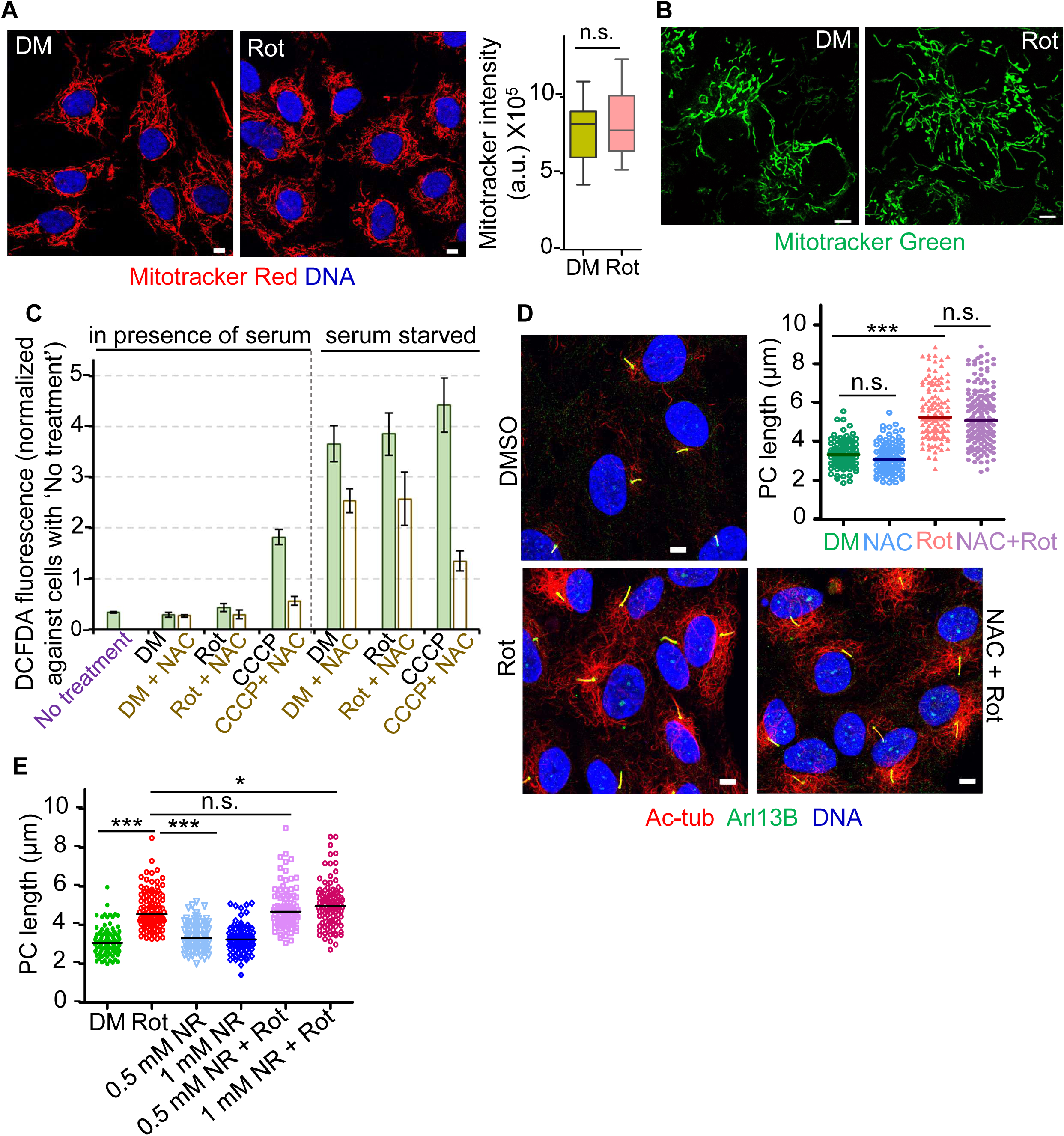
Rotenone-induced ciliary elonation is independent of mitochondrial depolarisation, ROS accumulation, and NAD⁺ availability (A) Serum starved RPE1 cells treated with DMSO and rotenone were incubated with MitoTracker Red and were fixed. Representative comfocal images are shown. Corrected total cell fluorescence (CTCF) of MitoTracker Red signal per cell are determined using ImageJ software and plotted as box and whisker, which show no significant change in its intensity under these conditions. (B) Serum starved RPE1 cells treated with DMSO and rotenone, and incubated with MitoTracker Green were imaged live and representative confocal images are shown here to demonstrate no gross alteration in mitochondrial morphology, content or membrane potential in these cells under these conditions. (C) Intracellular ROS levels in asynchronously growing or 24 h serum starved RPE1 cells, with or without NAC pre-incubation and treated with rotenone, CCCP or control solvent DMSO were measured by DCFDA fluorescence intensities. Fold increase in DCFDA fluorescence in each of the samples compared to ‘no treatment’ are plotted as bars, where values represent mean ± S.D., n=3. (D) Serum-starved RPE1 cells were treated with rotenone in the presence or absence of 1 mM NAC. Representative confocal images show that NAC does not rescue rotenone-induced ciliary elongation or cytoplasmic microtubule hyperacetylation, while the dot plots of PC length quantify the significance of the former. (E) Serum starved RPE1 cells were supplemented with nicotinamide riboside (NR) at the indicated concentrations, and then were treated with rotenone or DMSO, fixed and stained for ciliary markers. PC length was determined for all these samples, and are shown as dot plots, which clearly demonstrate that co-treatment with NR fails to prevent rotenone-induced PC elongation, suggesting that the phenotype is not driven by NAD⁺ depletion.

As rotenone treatment is frequently associated with elevated ROS level, we next evaluated whether oxidative stress due to enhanced ROS contributed to the observed phenotypes. We measured cellular ROS levels in both proliferating (serum-supplemented) and serum-starved conditions by quantifying DCFDA fluorescence in treated live cells. We observed that serum starvation itself produced a significant increase in the cellular ROS level compared to cells growing in presence of serum, which is not permissive condition for robust PC assembly in these cells (Fig.3C). However, treating cells with rotenone did not further elevate ROS levels under either condition. In the same assay condition, CCCP, an uncoupler of mitochondrial oxidative phosphorylation expectedly increased ROS in these conditions[25], which were further reduced to basal level upon pre-treating cells with N-acetylcysteine (NAC), an antioxidant that serves as ROS scavenger (Fig. 3C). This observation suggests that brief exposure to rotenone is not sufficient to enhance cellular ROS level to an extent that can affect cytoskeletal dynamics. Anyway, we proceeded to further examine if pre-treating cells with NAC may attenuate the observed effect of rotenone treatment on microtubule hyperacetylation or PC elongation. Our results clearly show that NAC pre-treatment did not reverse the rotenone-induced elongation of PC axoneme, while the microtubule hyperacetylation also remained prominent (Fig. 3D). These data further demonstrate that the ciliary elongation and microtubule hyperacetylation observed in rotenone treated cells are independent of ROS.

Given that rotenone can indirectly decrease intracellular NAD⁺ pools while inhibiting mitochondrial complex I[3], we tested whether supplementing NAD⁺ using nicotinamide riboside (NR) at 0.5-1 mM could mitigate the PC elongation phenotype. We observed that pre-incubation with NR did not reverse the PC elongation phenotype due to rotenone treatment, as compared to rotenone alone (without pre-treatment with NR; Fig. 3E). These findings indicate that supplementing the medium with NAD⁺ likely replenished the attenuated NAD⁺ as seen in other studies, but replenishing NAD⁺ does not regulate the ciliary or cytoskeletal responses induced by 100 nM rotenone treatment for 4 h.

Together, these results demonstrate that an acute, low-dose rotenone exposure does not induce significant mitochondrial toxicity, and thus the observed alterations in microtubule hyperacetylation or PC axoneme length triggered by such rotenone treatment occur independently of any mitochondrial toxicity. It is consistent with a direct effect of rotenone on microtubule dynamics rather than mitochondrial dysfunction.

### Rotenone-induced ciliary elongation occurs independently of microtubule hyperacetylation and is accompanied by attenuated intraflagellar transport

Because rotenone treatment led to a robust increase in microtubule acetylation (Fig. 1), we next asked whether this post-translational modification is required for the observed increase in PC length. To test this, we depleted α-TAT1, the primary α-tubulin acetyltransferase, using a previously validated siRNA[26] and examined the acetylation status for both cytoplasmic and the ciliary axonemal microtubules. As expected, α-TAT1 knockdown largely eliminated acetylation on both microtubules as seen by immunoblotting of total cellular tubulin and immunofluorescence of both DMSO and rotenone treated cells (Fig. 4A-C). Surprisingly, despite the near-complete loss of tubulin acetylation, rotenone continued to induce significant PC elongation, as judged by Arl13B staining that specifically mark ciliary membrane (Fig. 4A-B). These results indicated that microtubule hyperacetylation is not required for rotenone-induced PC elongation, considering that the observed near-complete loss of Ac-tub staining in α-TAT1-depleted cells indicate loss of only one type of microtubule modification, and not the loss of whole microtubular axoneme. To further validate that, we next stained these cells using the antibody (GT335) that specifically recognises glutamylation at c-terminal tails of protofilaments of stable microtubules such as that are present in ciliary axoneme. This critical post-translational microtubule modification is carried out by Tubulin tyrosine ligase-like (TTLL) enzymes, mostly by TTLL6 in human cells, and is not dependent on α-TAT1[27, 28]. In siControl and α-TAT1–depleted cells stained with GT335 antibody, rotenone treatment resulted in a similar increase in PC length, compared to DMSO treated cells (Fig. 4D). The persistence of elongation under two independent labelling conditions indicates that rotenone-induced PC elongation is indeed an increase in its axoneme length and is independent of its microtubule acetylation.

**Figure 4.**
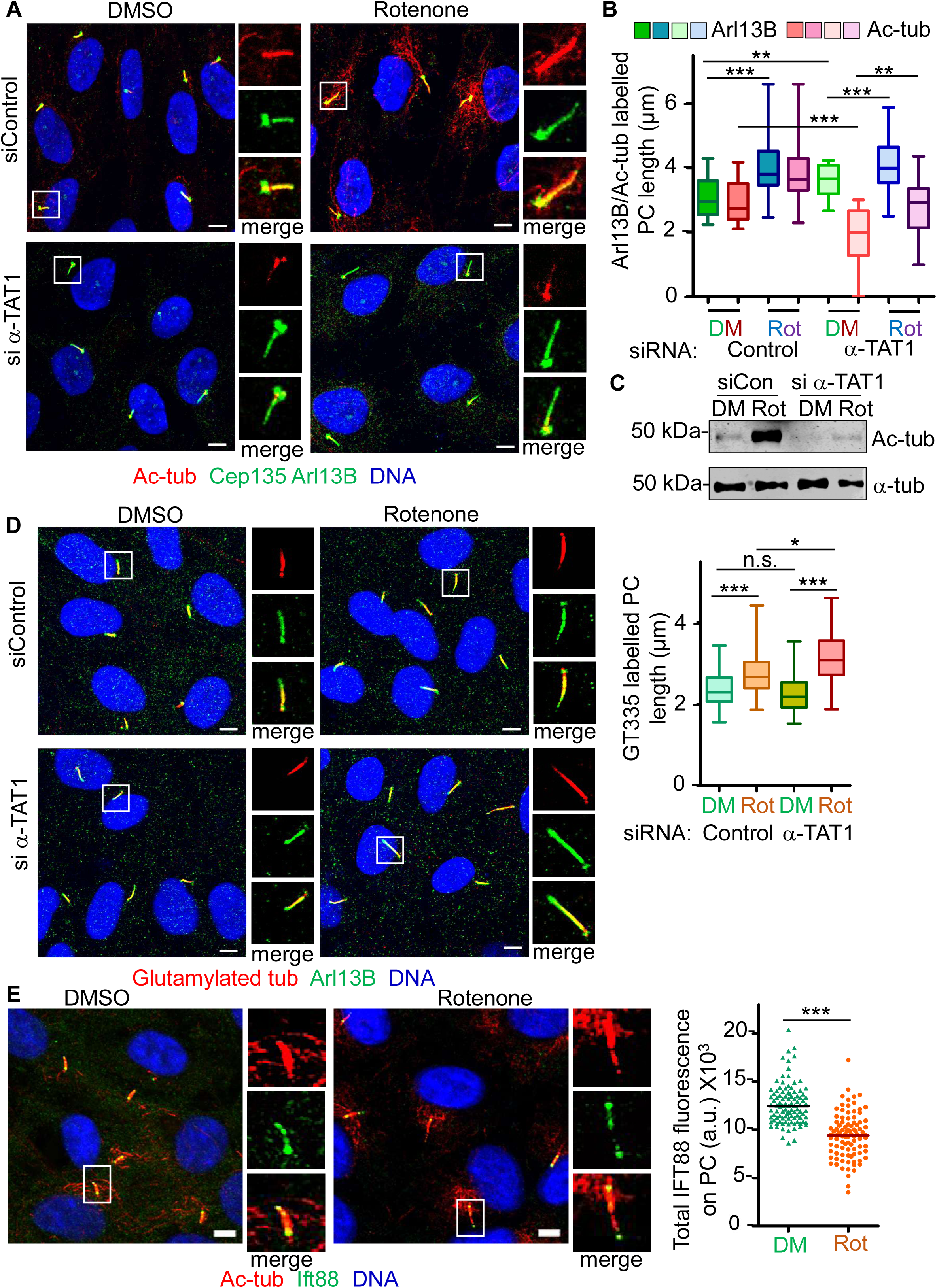
Rotenone-induced PC elongation is independent of α-TAT1-mediated microtubule acetylation (A) RPE1 cells transfected with control (siControl) or α-TAT1 (si α-TAT1) siRNAs were serum-starved for 24 h and treated with DMSO or 100 nM rotenone for 4 h. Representative confocal images of cells stained for Ac-tub and Arl13B are shown here, which show near-complete loss of acetylated microtubules in α-TAT1-depleted cells under both control and rotenone conditions. (B) Length of Ac-tub or and Arl13B stained PCs were determined separately in all four types of samples of A, and are shown as box and whisker plot. Rotenone significantly increases Arl13B-positive PC length in both control and α-TAT1 knockdown cells, demonstrating that rotenone-induced ciliary elongation occurs independently of α-TAT1–mediated tubulin acetylation. (C) Validation of α-TAT1 knockdown in these samples of A, as judged by the acetylated tubulin level in whole cell extracts, where α-tubulin was used as control for total tubulin content of those cell extracts. (D) Serum-starved siControl and si α-TAT1 cells treated with DMSO or rotenone were stained for glutamylated tubulin and Arl13B, and the representative confocal images were shown. The length of PC stained for glutamylated tubulin were determined and shown as box and whisker plot, demonstrating that rotenone still induces a significant increase in ciliary axoneme length under conditions where acetylation is abolished, supporting an actual structural elongation independent of Ac-tub labelling. (E) Representative confocal images of cells treated with DMSO or rotenone were stained for IFT88 and Ac-tub. Background-corrected total fluorescence intensities of IFT88 along Ac-tub-positive PCs are shown as dot plot, which demonstrate that rotenone treatment results in a significant reduction of IFT88 signal along the ciliary axoneme, suggesting impaired intraflagellar transport in elongated cilia.

We suspected that the drastic increase in the axoneme length could be due to an enhanced intraflagellar transport (IFT) in rotenone treated cells. Therefore, we examined the distribution of IFT88, the critical component of the IFT-B complex responsible for anterograde cargo movement. Immunostaining revealed detectable IFT88 along the axoneme in both DMSO and rotenone conditions. However, the total IFT88 signal along the PC was significantly reduced in rotenone-treated cells (Fig. 4E). This result suggests an aberrant anterograde IFT process upon rotenone treatment with the conserved amount of IFT88 molecules that show altered distribution along the PC, implying compromised transport activity. Alternatively, the kinetics of the anterograde IFT process in rotenone treated cells vary significantly than that in control cells, which may not be critically dependent on IFT88 activity. While deeper investigation of the altered IFT due to rotenone treatment is required to resolve the issue, it is beyond the scope of this present study. Nonetheless, our results clearly demonstrate that the observed increase in cytoplasmic microtubule acetylation and PC length are two separate outcomes of brief rotenone treatment in quiescent cells, which are related but not dependent on each other. Moreover, despite the increased length, rotenone-treated PC exhibit aberrant IFT process that may explain functional impairment reflected in transducing SHH signaling as seen earlier (Fig. 2).

### Rotenone treatment increases the soluble tubulin pool via microtubule depolymerization, which contributed to elongation of ciliary axoneme

Rotenone has been previously reported to promote microtubule depolymerisation[6, 29]. However, our study demonstrates that brief exposure to low concentration of rotenone increased hyperacetylation of cytoplasmic and ciliary microtubules and increased axoneme length, which are likely associated with enhanced stability of microtubules. We next sought to resolve this apparently conflicting aspect of how rotenone affects microtubule dynamics. We utilized high resolution confocal microscopy to examine the microtubule network stained by anti-α-tubulin antibody in rotenone treated serum starved RPE1 cells, and compared with DMSO treated cells. Additionally, we also treated cells separately with 1 μM taxol and 1 μM nocodazole for 2 h that promote irreversible tubulin polymerization or robust depolymerization of microtubules by inhibiting tubulin polymerization respectively. Expectedly, taxol treatment generated thick microtubule bundles, while nocodazole produced collapse of the microtubule network and near-complete loss of microtubule filaments that were seen in control cells (Fig. 5A-B). Importantly, rotenone-treated cells displayed significant loss of microtubule filaments and diffuse α-tubulin distribution throughout the cells, indicating microtubule depolymerization is promoted. Although the extent of microtubule depolymerization upon rotenone treatment appears to be lesser than that is seen in nocodazole-treated cells, microtubule network looks porous due to local microtubule depolymerization rather than catastrophic loss in rotenone-treated cells. Ultrastructural studies demonstrated that the lys40 of α-tubulin that is acetylated resides luminally in the microtubule filament with limited access to this site by the acetyl transferase enzymes such as α-TAT1 [30]. However, the slow kinetics of α-TAT1 also contributes to the overall rate, thereby suggesting that microtubules with slow dynamics are better substrates[31]. Accordingly, taxol treated cells show hyperacetylation of the stable microtubule bundles (Fig. 5C). However, perhaps it is not surprising that rotenone treatment facilitates hyperacetylation of cytoplasmic microtubules since due to porous nature of microtubule network in rotenone treated cells, the access of acetyl transferase to the site of acetylation is highly favoured (Fig. 5C)[32, 33]. In contrast, acetylated microtubules were nearly absent nocodazole-treated cells, likely owing to extensive microtubule depolymerisation. Notably, the thickness of the acetylated microtubules is significantly less in rotenone treated cells compared to that in taxol treated cells (Fig. 5C).

**Figure 5.**
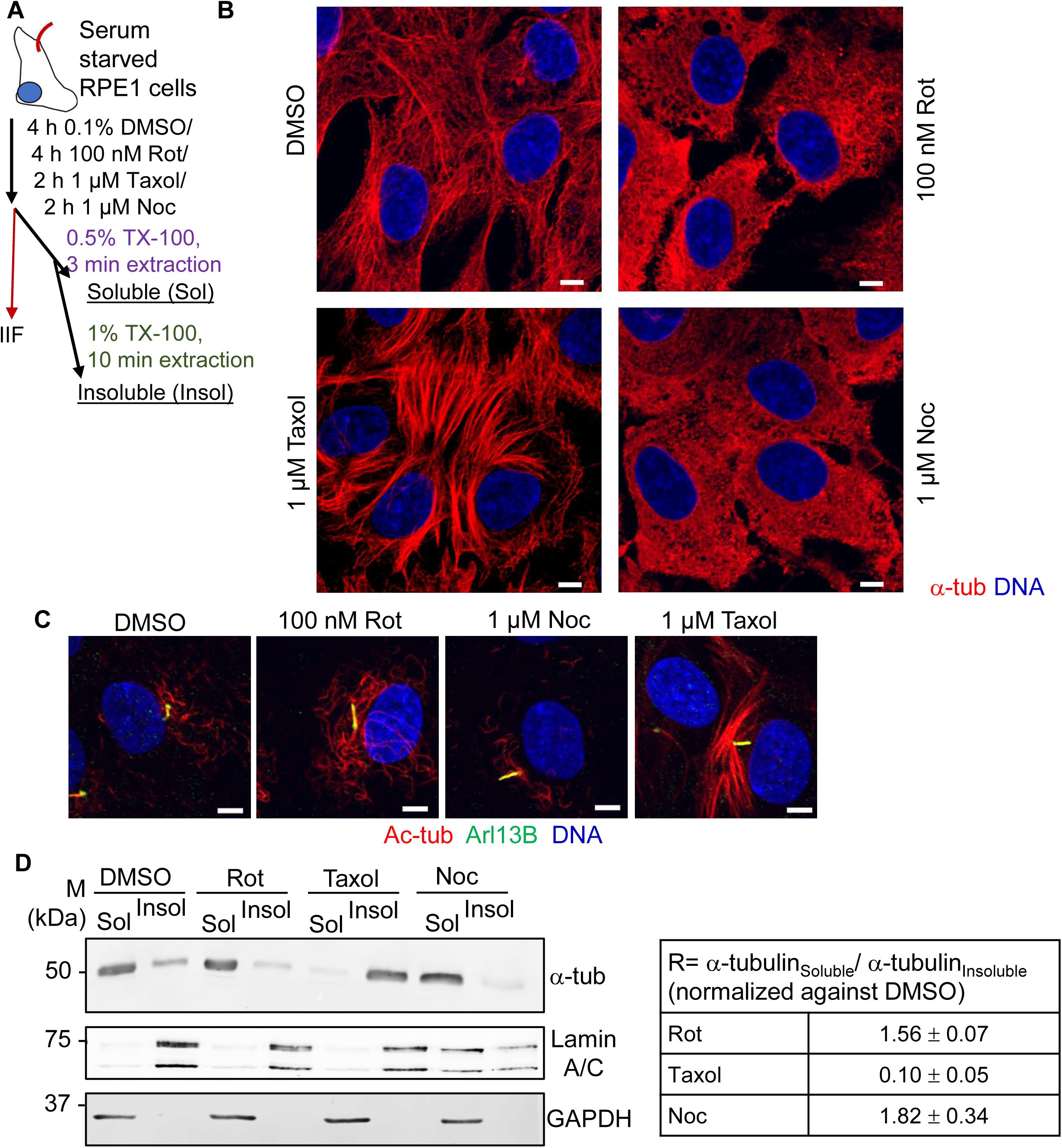
Rotenone increases the soluble tubulin pool while maintaining microtubule hyperacetylation, indicating a distinct mode of cytoskeletal remodelling. (A) Schematic of cellular fractionation into soluble cytosolic proteins and insoluble cytoskeleton and membrane bound proteins, the workflow used to separate soluble cytosolic and insoluble polymerised tubulin pools. (B) Serum starved RPE1 cells were treated with rotenone, taxol nocodazole or control solvent DMSO, fixed and stained for α-tubulin to examine cytoskeletal microtubule network. Representative images from high-resolution confocal microscopy are shown here, which show that compared to control cells, near-complete disruption of microtubule network in nocodazole treated cells, while rotenone also caused disruption of the cytoskeletal network, though in lesser extent than nocodazole. In contrast, taxol produces thick bundled microtubules. (C) Similarly treated cells were stained for Ac-tub and high-resolution confocal microscopy images are shown. (D) Immunoblot analysis of α-tubulin in soluble and insoluble fractions following treatment with the indicated small molecules. Immunoblotting using GAPDH and Lamin A/C antibodies validated the fractionation into soluble cytosolic proteins and insoluble cytoskeleton. The background corrected band intensities of α-tubulin were determined for each sample, and the ratio of α-tubulin in soluble and insoluble fraction are presented in the table, where values represent mean ± S.D., n=3. This data show that rotenone increases the soluble tubulin fraction to levels comparable to those of nocodazole, whereas taxol decreases the soluble pool, consistent with microtubule stabilisation.

Next, we analyzed the relative abundance of tubulin in soluble and insoluble cellular fractions following the protocol of an earlier study [13], where those crude fractions indicated unpolymerized and polymerized tubulin respectively. Like the previous experiment, we treated serum starved RPE1 cells briefly with taxol and nocodazole in addition to rotenone and DMSO, and extracted Triton X-100 soluble proteins that leaves insoluble cytoskeletal complex (Fig.5A). Immunoblotting of these two fractions show expected distribution of GAPDH in soluble fraction and nuclear membrane associated intermediate filament component Lamin A/C in the insoluble fractions in all four treatments. As expected, nocodazole treatment caused an almost two-fold increase in the soluble tubulin compared to polymerized tubulin, while taxol strongly reduced the soluble tubulin pool, consistent with its microtubule stabilising effect (Fig. 5D). Importantly, rotenone caused a significant increase in the soluble tubulin compared to the DMSO treatment, yielding a soluble-to-insoluble ratio to roughly 1.5-fold, suggesting that rotenone exerts a microtubule-depolymerizing effect that is similar to that of the destabilizing effect of nocodazole, al bait of moderately lesser extent. An earlier study reported that rotenone treatment Increase in soluble tubulin in mouse dopaminergic (DA) neurons due to microtubule depolymerizing effect. That was one of the reasons, and not mitochondrial complex I inhibition, for rotenone induced toxicity of mouse dopamine neurons [7].

Notably, soluble cytoplasmic tubulin molecules usually concentrates near centrosomes[34], and an increase in this soluble tubulin pool promotes drastic elongation of ciliary axoneme [13]. Based on this information and our observations, we propose that the observed drastic increase in PC length due to brief treatment of rotenone occurs due to increase in soluble cytoplasmic tubulin and simultaneously increased stabilization and flexibility of microtubular filaments due to hyperacetylation in these non-mitotic cells. Since a similar mechanism of increase in soluble tubulin happens in dopaminergic mouse neurons, it is tempting to suggest that brief exposure to low concentration of rotenone may impair primary cilia structure, and the signaling pathways via these cilia in non-mitotic dopaminergic neurons, which may contribute to the pathology of Parkinson’s disease and other neuronal ciliopathies.

## Discussion

Rotenone is widely used as a mitochondrial complex I inhibitor and as an experimental inducer of Parkinson’s disease-like phenotypes, with most studies focusing on its roles in mitochondrial dysfunction, oxidative stress, followed by dopaminergic neuronal death. It is fascinating that primary cilia length was shown to be altered with impaired SHH activity in dopaminergic neurons derived from PD patients and in mice model of PD[15]. In this study, we discover a new mode of rotenone toxicity that depends on rotenone’s effect of microtubule network and dynamics, and independent of rotenone’s impact on mitochondrial membrane potential, increase of cellular ROS or NAD⁺ depletion. Our observations are consistent with previous studies that demonstrated rotenone’s impact on cytoskeletal dynamics. However, this study first show that even short-term, low-dose exposure to rotenone results in drastic PC elongation in quiescent cells that are not undergoing proliferation.

Our study demonstrates that acute rotenone exposure induces a rapid and striking elongation of primary cilia in cell types like RPE1 and U87-MG glioblastoma cells that at least assemble PC. These elongated cilia were consistently accompanied by hyperacetylation of cytoplasmic microtubules, a modification often associated with increased microtubule stability. Such hyperacetylation in rotenone treated cells is conserved across various cells irrespective of PC assembly such as neuroblastoma SHSY-5Y cells or rat PC12 cells[22]. On the other hand, long-term rotenone treatment induced PC elongation independent of cell cycle regulation and cellular ATP level but dependent on mitochondrial ROS level in proliferating IMCD3 and RPE1 cells[21]. However, despite their increased length, rotenone-treated PC were compromised in transducing SHH signaling as these cells fail to show uniform Smo distribution along PC and transcriptional increase in canonical SHH target genes upon SAG activation. This uncoupling of ciliary length from signalling capacity highlights that even subtle perturbations in microtubule dynamics lead to dysfunctional PC.

However, our mechanistic investigation revealed that microtubule acetylation is not required for the rotenone-induced elongation phenotype. Depletion of α-TAT1 eliminated acetylation from both the cytoplasm and the axonemal microtubules, although the axoneme remained elongated in rotenone treated cells, which was further confirmed by examining the glutamylation status of axonemal microtubules. Thus, rotenone treatment affects specific PTMs on ciliary microtubules. Rotenone treatment also affected IFT88 levels along the PC, reiterating that aberrant elongation of PC is coupled with impaired IFT, which is one plausible explanation for the attenuated Smo level on PC upon SAG-mediated activation of SHH signalling. Although Smo can enter the cilium independently of IFT, its proper distribution along the axoneme and the stabilisation of its signalling-competent state require functional IFT components, such as IFT88 [35]. Thus, decreased ciliary IFT88 level may limit the cilium’s ability to traffic signalling cargo, resulting in a structurally enlarged but functionally “silent” cilium.

Since rotenone is classically linked to oxidative stress, we rigorously tested whether ROS contributes to these effects. Across multiple assays, including DCFDA quantification, antioxidant rescue with NAC, and NAD⁺ supplementation with NR, we found that the ciliary and cytoskeletal changes persist in the absence of elevated ROS or NAD⁺ depletion. Earlier studies suggested that rotenone-induced PC elongation is not dependent on its ability to inhibit mitochondrial complex I[21].

Instead, our data support a model in which rotenone directly or indirectly perturbs microtubule polymerisation-depolymerisation dynamics. Consistent with other studies showing that rotenone can bind to tubulin and promote microtubule depolymerization[6], our data revealed a marked increase in the soluble tubulin pool upon rotenone exposure, comparable to that of nocodazole, despite simultaneous hyperacetylation of the microtubules. Thus, rotenone treatment had likely stabilized the microtubules via hyperacetylation of the polymers, suggesting that acetylation may represent a compensatory or stress-induced modification rather than a purely stabilising event. Together, these findings suggest a model in which rotenone shifts the balance of microtubule assembly toward increased turnover, resulting in a cytoskeletal environment that promotes ciliary elongation. Such possibilities are consistent with few previous studies on rotenone’s impact on cytoskeletal dynamics [2, 6, 7, 29], although these studies did not consider all the aspects to explain altered PC structure and function due to such short-term, low-dose exposure to rotenone.

The implications of this work extend beyond in vitro observations. PCs are increasingly recognised as critical regulators of neuronal development, synaptic plasticity, neurogenesis, and adult tissue homeostasis[15, 16, 18, 36, 37]. Disruption of ciliary signalling, including SHH pathway dysfunction, is linked to a broad spectrum of neurological and developmental disorders. Our findings suggest that rotenone-induced ciliary dysfunction in quiescent cells may contribute to neurodegenerative processes not only through mitochondrial stress but also by compromising cilia-dependent signalling pathways essential for neuronal health. This dual impact, a metabolic insult combined with disruption of primary cilia mediated signaling, may help explain why rotenone treatment models reproduce complex aspects of Parkinson’s pathology that cannot be attributed solely to loss of mitochondrial function[2, 5, 7].

Several important questions remain. How does rotenone simultaneously increase microtubule acetylation yet increase the soluble tubulin pool? How these soluble tubulin molecules enter cilia and participate in axoneme elongation when the IFT-B process appears compromised? An earlier study in *Chlamydomonas* showed the contribution of anterograde IFT process in facilitating soluble tubulin to increase cilia length although the possibility of diffusion of tubulin molecules inside cilia was not completely neglected[38]. Alternatively, is the reduction in IFT88 an indirect consequence of altered microtubule dynamics? Finally, the question remains if similar ciliary defects occur in neurons exposed to rotenone in vivo, particularly in dopaminergic neurons that are known to rely heavily on ciliary signalling? While these critical questions are beyond the scope of the present study, we will look forward to address them in near future.

In summary, our study reveals a novel dimension of rotenone toxicity involving cytoskeletal remodelling and primary cilia dysfunction, distinct from its canonical mitochondrial effects or perturbation of cell cycle in quiescent cells. These findings open new avenues for investigating how environmental toxins perturb ciliary architecture and signalling and lay the groundwork for future studies aimed at understanding the contribution of ciliary dysregulation to neurodegenerative disease.

## Materials and methods

### Cell culture

RPE1 (ATCC) and U-87 MG (ATCC) cells were cultured in DMEM with 10% FBS (Himedia) and 1% Penicillin-Streptomycin (Thermo) at 37°C in 5% CO2. For serum starvation, cells were incubated in serum-free DMEM medium supplemented with 1% Penicillin-Streptomycin for the indicated time. Most commonly, 24 h serum starved cells were treated with rotenone (Sigma) for indicated time. For N-acetylcysteine (NAC, Sigma) and nicotinamide riboside (NR, MedChemExpress) treatments, serum starved cells were subjected to indicated final concentration of NAC for 1 h or NR for 24 h, followed by treatment with 100 nM rotenone for 4 h. Cells were treated with nocodazole or taxol (both from Sigma) at the final concentration of 1 μM. Rotenone, nocodazole and taxol were dissolved in cell culture grade DMSO (Sigma), whereas NAC and NR were dissolved in deionized water. To stimulate SHH activation, 500 nM Smoothened agonist (SAG; MedChemExpress) was used for the mentioned duration, followed by rotenone treatment. To measure the cellular Reactive Oxygen Species (ROS), 10,000 RPE1 cells were seeded in a 96-well cell culture plate and grown for 24 h. Then, the cells were either serum-starved or maintained in growing media for the next 24 hours. Next, cells were either left untreated or treated with 1 mM NAC for 1 h, followed by treatment with rotenone (4 h), CCCP (1 h), or control solvent (DMSO) (4 h). For the last 30 min of the treatment, cells were incubated with 20 µM 2’,7’-dichlorodihydrofluorescein diacetate (DCFDA, Sigma), and after washing with warm PBS (Gibco), DCF fluorescence was recorded using a BioTek Synergy microplate reader (Agilent) following manufacturer’s instruction. To assess cell viability, 10,000 RPE1 cells were seeded in a 96-well transparent cell culture plate and grown for 24 h before changing the media to complete media (DMEM with 10% FBS, 1% Penicillin-Streptomycin) or serum-free media. The MTT reagent was added at a final concentration of 0.5 mg/mL, and the cells were incubated for 1.5 h in the cell culture incubator at 37°C and 5% CO2. Next, the media was removed, formazan precipitate was dissolved in an equal volume of DMSO, and OD_570_ was measured using a BioTek Synergy microplate reader following manufacturer’s instruction.

### siRNA Transfection

Previously validated siRNA against α-TAT1 (5′-AACCGCCATGTTGTTTATATT-3) was purchased from Qiagen [22, 26], while a silencer negative control (Ambion, Thermo [39]) was used as control. Transfection of siRNAs at a final concentration of 10 nM was performed with Lipofectamine RNAiMAX (Thermo) according to the manufacturer’s instructions.

### Cytology

Cells were fixed in either 4% paraformaldehyde-0.2% TritonX-100 for 10 mins at room temperature or chilled methanol at -20 °C for 10 mins. Primary antibodies used here were: mouse anti-γ tubulin (1:200, CST); rabbit anti-Arl13B (1:250, Proteintech); mouse anti-acetylated α-tubulin (Ac-tub; 1:1000, Sigma), mouse anti-Smoothened (Smo; 1:200, Santa Cruz), mouse anti-Glutamylated tubulin (GT335; 1:5000, Adipogen, kind gift from Dr Sudarshan Gadadhar), rabbit anti-IFT88 (1:500, Proteintech; kind gift from Dr Swapnil Shinde). Secondary antibodies were donkey anti-rabbit or donkey anti-mouse, tagged with either AlexaFluor 488 or AlexaFluor 568 (1:1000; Invitrogen). Hoechst33358 (1 μg/ml; Sigma) was used to stain nuclear DNA. To assess mitochondrial health, cells were incubated with 100 nM MitotrackerRed (Invitrogen) for 45 minutes and then fixed with 4% Paraformaldehyde (EMS). Live imaging of mitochondria was performed using MitoTracker Green (Invitrogen; kind gift from Dr Piyali Mukherjee) at a final concentration of 100 nM for 45 min. Images of fixed cells were acquired using a Zeiss Fluorescence microscope fitted with a 63X Plan Apo oil immersion objective (NA 1.4). A Leica Confocal laser scanning microscope, equipped with a 63X Plan Apo oil immersion objective (NA 1.4), was used to acquire images of fixed and live cells. The length of the PC (in µm), stained for Ac-tub, GT335 or Arl13B was determined by the associated software packages (Zen blue3.3 for Zeiss or LasX for Leica).

### Immunoblotting

The BCA protein assay method (Pierce) was used to determine the total protein concentration of cell lysates prepared in a buffer containing 50 mM Tris-Cl, pH 8.0, 150 mM NaCl, and 1% NP-40, supplemented with protease inhibitors (Sigma). Antibodies for immunoblotting were: mouse anti-acetylated tubulin (1:5000; Sigma), mouse anti-α-Tubulin DM1A (1:5000; Sigma), mouse anti-GAPDH (1:5000; NOVUS), and rabbit anti-Lamin A/C (1:2000, ABclonal). Alexafluor680-conjugated donkey anti-rabbit and DyLight800-conjugated donkey anti-mouse (both 1:10000; Invitrogen) secondary antibodies were used for fluorescence scanning using the Odyssey-DLx infrared imaging system (Li-Cor). The background-corrected intensities of bands were determined using ImageStudio (Li-Cor) or ImageJ software.

### qRT-PCR

Total RNA from cells was isolated using TRIzol (Invitrogen) according to the manufacturer’s protocol, and dissolved in 20 μL of nuclease-free water. 2 µg DNase-treated total RNA was then reverse transcribed using iScript cDNA synthesis kit (Biorad). The cDNA was subsequently used for quantitative, real-time PCR (qPCR) using SsoAdvanced Universal SYBR Green Supermix (Bio-Rad) on a CFX Connect Real-Time System (Bio-Rad), where the mentioned oligos were used at a concentration of 500 nM each. qPCR primer-pair efficiency (E) of all sets of primers were calculated using manufacturer’s protocol (Biorad; E-values ranging from 90%-110% were considered as efficient primer pairs). The sequences of the primers (also used in previous study[40]): GLI1_Fwd: CCTGGATCGGATAGGTGGTC, GLI1_Rev: ACCTCTGGACTCTAGCTGGA; PTCH1_Fwd: GCCCAGTTCCCTTTCTACCT, PTCH1_Rev: CAACACCACGCTGATGAACA; HHIP_Fwd: TGCAAAATGTGAGCCAGCAT, HHIP_Rev: GGTCACTCTGCGGATGTTTC; SMO_Fwd: TGGCACACTTCCTTCAAAGC, SMO_Rev: TGAGGACAAAGGGGAGTGAC; β- ACTIN_Fwd: GCTCACCATGGATGATGATATCGC, β- ACTIN _Rev: ATAGGAATCCTTCTGACCCATGCC

### Statistical analysis

Each experiment was performed at least three times with 2-3 technical replicates wherever applicable. Either a One-way ANOVA or an unpaired Student’s T-test was conducted to determine the significance level. The *p-value *<*0.05, **p-value *<*0.01, and ***p-value *<*0.001 were considered statistically significant. n.s. indicates non-significant.

## Acknowledgement

This work is supported by Ramalingaswami fellowship to SM by the Department of Biotechnology (DBT), Govt. of India, and the core research grant from the Science and Engineering Research Board (SERB, presently ANRF), Govt. of India (CRG/2020/004042) to SM. PH and RM are respectively University Grants Commission (UGC) and DBT research fellows. Authors sincerely thank Drs Chandrama Mukherjee and Piyali Mukherjee of Presidency University; Sudarshan Gangadhar of BRIC-inStem, Swapnil R. Shindey of IIT Bombay, Karthigeyan Dhanasekaran of DBT-Regional Centre of Biotechnology (RCB) and Oishee Chakrabarti of Saha Institute of Nuclear Physics for reagents. Authors are thankful to the microscope facility of Institute of Health Sciences, Presidency University, and the high-resolution microscopy facility (Nikon) of Saha Institute of Nuclear Physics and Dr Kaushik Sengupta, the coordinator of that facility. During the preparation of this manuscript the authors used ChatGPT (https://chatgpt.com/) to improve the English in certain cases and not for drawing any scientific conclusions.

